# A framework for reparative CAR T engineering in the CNS

**DOI:** 10.64898/2026.02.09.703122

**Authors:** Rotem Shalita, Maya Ben Yehuda, Pavle Boskovic, Yasmin Frid, Yuri Kuznetsov, Michael Tsoory, Vyacheslav Kalchenko, Ori Brenner, Eyal David, Kfir Mazuz, Robbie Majzner, Jonathan Kipnis, Ido Amit

## Abstract

Chimeric antigen receptor (CAR) T cells have shown remarkable therapeutic promise in hematological malignancies and, more recently, in autoimmunity. They also hold considerable potential for neurodegenerative diseases and CNS injury, where the therapeutic objective shifts from cell depletion to immune modulation and tissue repair. Using ischemic stroke as proof-of-concept, we engineered MOG-targeting CAR T cells to dissect how distinct CAR designs shape the CNS microenvironment. CD4/CD8 CAR T cells (a mixture of CD4⁺ and CD8⁺ subsets) proliferated robustly and efficiently infiltrated the ischemic hemisphere, but induced broad immune recruitment and exacerbated neuroinflammation. In contrast, CD4⁺ restricted CAR T cells markedly reduced immune infiltration, reprogrammed microglia, and minimized inflammatory activation. We engineered CD4⁺ CAR T cells to secrete brain-derived neurotrophic factor (BDNF) to determine whether they could be redirected toward a reparative, non-cytotoxic phenotype. CD4⁺ BDNF-CAR T cells further attenuated inflammation, reduced immune infiltration, and promoted the expansion of regulatory T cells. CD4⁺ BDNF-CAR T treated mice showed significantly improved gait performance following stroke. Together, these findings establish a cellular framework and outline principles for engineering reparative CAR T platforms for neurological diseases.

## Introduction

Chimeric Antigen Receptor (CAR) T cell therapy represents a rapidly expanding frontier in therapeutic development for neurological disorders, with growing success in both autoimmune neurologic diseases^1,2^ and malignant CNS tumors^3–5^. Compared with monoclonal antibodies, CAR T cells exhibit the capacity to actively traffic into and proliferate in tissues including the CNS^6,7^, making them an appealing platform for neurological applications. This opens substantial opportunities to repurpose CAR T cells for neurodegenerative diseases and CNS injury, conditions in which the goal is not cell elimination, but immunomodulation and promotion of tissue repair^8^. CAR T cells enable a targeted strategy to ameliorate immune dysfunction by selectively reshaping immune responses^9^.

Previous studies demonstrated that endogenous T cells play active and protective roles in neurodegeneration and CNS injury^10–18^. In particular, recent work on spinal cord injury has revealed clonal expansion of injury-associated T cells in both mice and humans, with CD4⁺ T cell clones recognizing self-antigens derived from myelin and neuronal proteins^10^. Transient reconstitution of these autoreactive T cells conferred neuroprotection by modulating myeloid cell states, providing direct evidence that T cells can actively promote repair in the injured CNS rather than exacerbate damage^10,13,14,17^. CD4⁺ T cells are particularly attractive for CAR T applications in the CNS, given their central role in coordinating immune responses, shaping myeloid cell behavior, and supporting tissue repair^10–14,17^. At the same time, emerging evidence suggests that CD8⁺ T cells can play a protective role in neurodegenerative settings by interacting with microglia^19–21^. Thus, limiting therapeutic design to CD4⁺ compartments may overlook key protective functions provided by CD8⁺ T cells. A systematic evaluation of both CD4⁺ and CD8⁺ CAR T cell programs is required to engineer an optimized therapeutic approach for T cell mediated immune modulation in the CNS.

The first aspect of immune modulation involves the delivery of therapeutic molecules. Cellular therapies are capable of sensing and integrating complex physiological cues to enable precise, context-dependent cargo release^22^. Recent studies in solid tumors and autoimmune conditions have shown that CAR T cells can be engineered to secrete a broad range of immunomodulatory factors directly within target tissues: these engineered cells can deliver pro-inflammatory cytokines^23–31^, anti-inflammatory factors^32,33^, antibodies that prevent CAR T cell exhaustion^34^ or act as bispecific cell engagers^35,36^, soluble receptor decoys that neutralize inflammatory cytokines^32^ or block tumor-mediated suppression^37^, and pro-apoptotic factors^38^ that boost tumor cell death. Precise tuning of these payloads allows CAR T cells to actively reprogram the local microenvironment while simultaneously enhancing their own cytotoxicity, persistence, and expansion.

Building on these advances, similar principles can be applied to neurodegenerative diseases and CNS injury, where CAR T cells could be programmed to deliver neurotrophic, anti-inflammatory, or pro-repair factors to support neuronal survival and restore immune balance. However, the optimal design principles for inducing beneficial immune modulation remain undefined. Current T cell mediated cargo-delivery strategies are limited, with only a few studies using CD4⁺ T cells or CAR regulatory T cells (Tregs) engineered to secrete brain-derived neurotrophic factor (BDNF) as a neuroprotective factor in neurodegenerative conditions^39,40^. Additionally, how BDNF-secreting CD4 T cells reprogram the immune microenvironment remains undefined. Careful evaluation is needed to understand how CAR T cells armored with distinct neurotrophic or immunomodulatory cargos influence not only the surrounding tissue environment but also CAR T cell phenotypes.

The second facet of immune modulation is recruiting and reprogramming other immune populations to support tissue repair. In neurodegeneration and CNS injury, immune responses are increasingly understood to be multifaceted^41^: infiltrating lymphocytes and CNS-resident microglia can exacerbate acute inflammation and tissue damage^42,43^, yet they also play essential roles in resolving inflammation and clearing debris^10,12–15,44,45^. It remains unclear which engineering strategies are best suited to recruit reparative immune subsets, reset inflammation, and shift microglia toward a disease-associated microglia (DAM) neuroprotective state^45^. This is especially critical in the CNS, where traditional CAR T cell therapies have been associated with neurotoxicity, including immune effector cell-associated neurotoxicity syndrome (ICANS)^46^. These risks highlight the need for rational design principles tailored specifically to neurological indications, where we can leverage the natural capacity of T cells to penetrate the CNS, secrete cytokines, and reprogram the local environment, while minimizing the cytotoxic response.

Using ischemic stroke as a proof of concept and building on evidence that self-reactive, myelin-recognizing T cells can exert neuroprotective functions in CNS injury^10,13,14^, we engineered MOG-targeting CAR T cells. We applied single-cell multiomics to analyze how CAR T cells interact with a damaged CNS environment to map both therapeutic and detrimental CAR T responses in the injury niche. We compared different CAR T approaches: combined CD4⁺ and CD8⁺ CAR T cells, CD4⁺-restricted CAR T cells, and BDNF secreting CAR T cells. This analysis revealed how the different approaches modulate immune cell infiltration, CNS cellular and molecular profiles, and CAR T cell phenotypes. Together, we outlined core engineering principles for achieving precise, CNS-specific CAR T cell activity and leveraged these insights to develop CAR T cell therapy for ischemic stroke that reprograms the CNS microenvironment in a hemisphere-specific manner and improves functional recovery. These findings establish a therapeutic framework for safely translating CAR T approaches into neurodegenerative disease and CNS injury.

## Results

### Cerebral ischemia induces robust immune alterations that underlie gait deficits

We first established and validated the ischemic injury model using the middle cerebral artery occlusion (MCAO) paradigm, as previously described^47,48^. To confirm successful occlusion and reperfusion, we applied transcranial optical vascular imaging (TOVI)^49^, an *in vivo* imaging technique that integrates laser speckle imaging and fluorescent angiography to provide high-resolution assessment of cortical blood flow (Figure 1A). TOVI revealed clear reductions in perfusion in the right hemisphere during occlusion and restoration of flow following removal of the filament (Figure 1A). Histological analysis further confirmed unilateral injury, as evidenced by red (eosinophilic) neurons and vacuolation in the right septal nuclei and caudate-putamen, consistent with focal ischemic and neuronal damage (Figures 1B, S1A). To characterize the immune response, we performed single-cell RNA sequencing (scRNA-seq) of immune cells isolated from the ischemic right and non-damaged left hemispheres 14 days post-MCAO, identifying diverse infiltrating and resident populations including T cells, NK cells, microglia, monocytes, monocyte-derived macrophages, dendritic cells, and B cells (Figures 1C, S1B). Mice that underwent MCAO exhibited broad immune remodeling relative to controls, characterized by the infiltration of peripheral immune cells and state changes in resident microglia (Figure S1C), consistent with previous reports^44^. Behavioral assessment of gait performance using the CatWalk system^50,51^ was performed to assess motor function. To account for heterogeneity in stroke severity, we separated mice that underwent MCAO into “deficit” and “non-deficit” groups based on early gait performance following surgery. Non-deficit animals represented ∼25% of mice that underwent MCAO but did not show reductions in speed, stride length, or swing speed 2 days after surgery, likely reflecting incomplete occlusion. This classification was further validated by blinded histopathology reports, indicating no detectable injury or abnormality in their brain tissue (data not shown). Finally, motor assessment of deficit-bearing mice demonstrated clear gait impairment, evident at 2 days and continuing through 1 week after MCAO, relative to their pre-surgery baselines, with reduced speed, increased run duration, decreased swing speed, and shorter stride length compared to sham controls (Figures 1D-G, S1D-G). These deficits were not confined to the contralateral side of the body, as would typically be expected following a unilateral lesion in humans, but were observed bilaterally in our MCAO model, consistent with previous reports^52^ and potentially reflecting bilateral compensatory gait adjustments. Together, these vascular, histological, molecular, and behavioral analyses comprehensively validate the MCAO model, providing a foundation for evaluating the therapeutic potential of CAR T cells in the post-ischemic immune environment.

**Figure 1.**
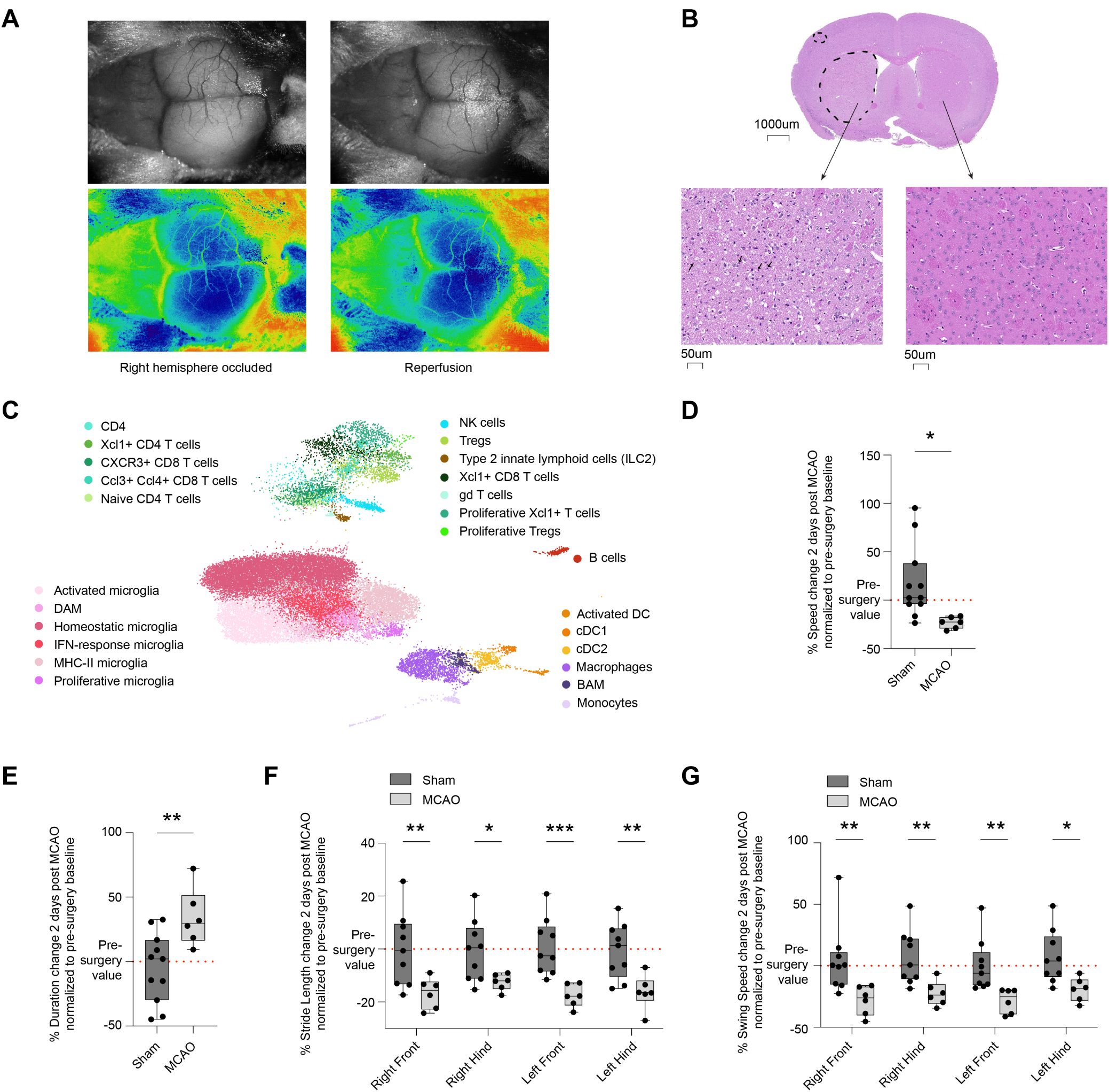
Cerebral ischemia induces robust inflammatory responses, infiltration of peripheral leukocytes, and results in gait deficits. (A) TOVI imaging in a living mouse, showing the right hemisphere occluded with a filament inserted to the middle cerebral artery (MCA), followed by filament withdrawal and reperfusion of the right hemisphere. (B) H&E staining of a representative mouse brain 5 days post 60 minute occlusion of the MCA. The cerebrum was cut into 4 slices which were laid on the caudal aspect. Microscopic changes were identified in the 3 cranial sections in a unilateral distribution. The section shown is the septal nuclei and caudate-putamen which had the most widespread changes. A cluster of red neurons with fine vacuolation of the neuropil is present in the dorsal cortex. Isolated red neurons are found in a large area of blue neurons in the lateral cortex and in the ventral cortex. The most obvious changes affect the entire caudate putamen in which there is widespread vacuolation and scattered red neurons. Vacuoles vary from minute to ∼20µ. (C) UMAP visualization of scVI embedding on 29,892 single cells obtained from the brains of mice that underwent MCAO and controls. Color represents the different major cell-type lineages obtained by graph-based clustering and annotated by conventional markers. (D) Boxplot showing percent speed change 2 days post MCAO surgery, normalized to pre-surgery baselines, and depicting mice that underwent MCAO with gait deficits and controls. Statistical significance was determined using an unpaired t-test. ∗p≤0.05. (E) Boxplot showing percent duration change 2 days post MCAO surgery, normalized to pre-surgery baselines, and depicting mice that underwent MCAO with gait deficits and controls. Statistical significance was determined using an unpaired t-test. ∗∗p≤0.01. (F) Boxplot showing percent stride length change 2 days post MCAO surgery, normalized to pre-surgery baselines, and depicting mice that underwent MCAO with gait deficits and controls. Statistical significance was determined using an ordinary two-way ANOVA with Sidak’s multiple comparisons test. ∗∗∗p≤0.001, ∗∗p≤0.01, ∗p≤0.05. (G) Boxplot showing percent swing speed change 2 days post MCAO surgery, normalized to pre-surgery baselines, and depicting mice that underwent MCAO with gait deficits and controls. Statistical significance was determined using an ordinary two-way ANOVA with Sidak’s multiple comparisons test. ∗∗p≤0.01, ∗p≤0.05.

### CAR T cells enable spatially precise immune reprogramming in the post-ischemic CNS

Having established a reference state of post-ischemic neuroinflammation, we next engineered CAR T cell constructs to evaluate their reparative potential within the injured CNS. As a CNS-restricted antigen, we selected myelin oligodendrocyte glycoprotein (MOG) as the CAR target (Figure 2A). We designed a second-generation anti-MOG (aMOG) CAR incorporating an scFv derived from an anti-MOG antibody^53^, a mouse CD28 co-stimulatory domain, a mouse CD3ζ signaling domain, and an EGFP reporter within a retroviral backbone (Figure 2B). An additional variant incorporated mouse CD8 hinge and 4-1BB cytoplasmic domains, followed by mouse CD3ζ signaling domain (Figure S2A). Antigen specificity was validated by IFN-γ secretion, which showed strong and selective activation in response to myelin stimulation (Figures 2C, S2B). The CD28-CD3ζ configuration produced the strongest antigen-dependent response (Figure S2B) and was selected for downstream *in vivo* studies.

**Figure 2.**
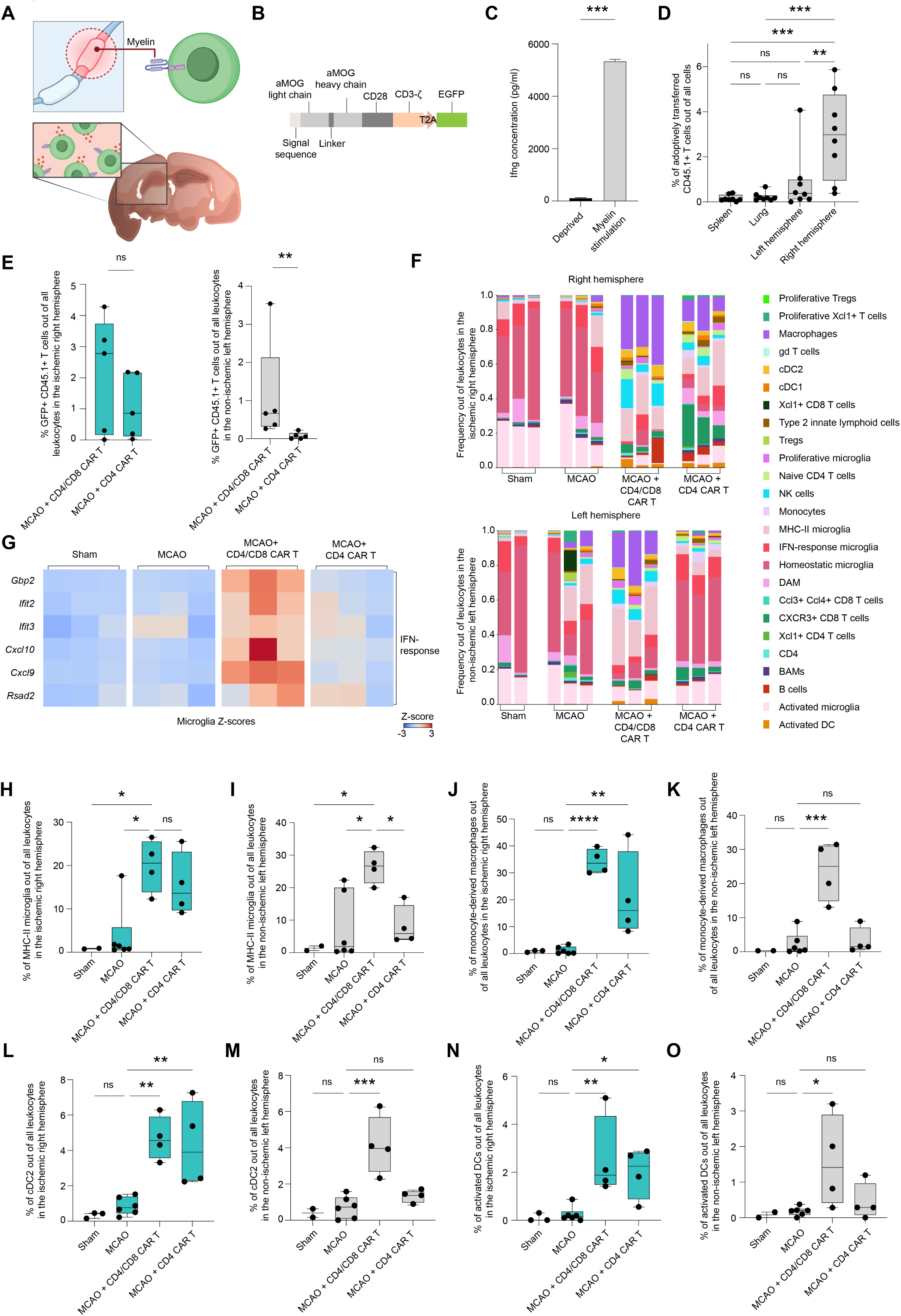
CAR T cells enable spatially precise immune reprogramming in the post-ischemic CNS. (A) Schematic illustration of aMOG CAR T cells and their homing to the damaged site. (B) Schematic description of aMOG CD28 CD3z CAR vector. (C) Mouse IFN-γ ELISA assay of aMOG CD4/CD8 CAR T cells, either unstimulated or stimulated with myelin. Statistical significance was determined using an unpaired t-test.. ∗∗∗p≤0.001. (D) Boxplot showing frequency of adoptively transferred CD45.1+ T cells out of all cells in the spleen, lung, left and right hemispheres. Statistical significance was determined using an ordinary one-way ANOVA with Tukey’s multiple comparisons test. ∗∗∗p≤0.001, ∗∗p≤0.01. (E) Boxplots showing frequencies of adoptively transferred GFP+ CD45.1+ T cells out of all cells in each hemisphere, for mice that underwent MCAO and were treated with either 10M CD4/CD8 CAR T or 10M CD4 CAR T. Statistical significance was determined using a two-tailed Mann-Whitney U test. ∗∗p≤0.01. (F) Bar plots showing the relative abundance of each cell population out of all leukocytes in the right ischemic hemisphere (top) and left non-ischemic hemisphere (bottom) of sham controls, and mice induced with MCAO who were either injected with PBS, 10M aMOG CD4/CD8 CAR T cells, or 10M aMOG CD4 CAR T cells immediately following MCAO. (G) Heatmap showing IFN-response gene expression of microglia in the brains of sham controls, and mice induced with MCAO who were either injected with PBS, 10M aMOG CD4/CD8 CAR T cells, or 10M aMOG CD4 CAR T cells immediately following MCAO (n = 12). The values are row-normalized z-scores of the pseudobulk gene expression. Each column represents the pooled average reads of a mouse. (H) Boxplot showing frequencies of MHC-II microglia out of all leukocytes in the ischemic right hemisphere of sham controls, and mice induced with MCAO who were either injected with PBS, 10M aMOG CD4/CD8 CAR T cells, or 10M aMOG CD4 CAR T cells immediately following MCAO. Statistical significance was determined using an ordinary one-way ANOVA with Tukey’s multiple comparisons test. ∗p≤0.05. (I) Boxplot showing frequencies of MHC-II microglia out of all leukocytes in the non-ischemic left hemisphere of sham controls, and mice induced with MCAO who were either injected with PBS, 10M aMOG CD4/CD8 CAR T cells, or 10M aMOG CD4 CAR T cells immediately following MCAO. Statistical significance was determined using an ordinary one-way ANOVA with Tukey’s multiple comparisons test. ∗p≤0.05. (J) Boxplot showing frequencies of monocyte-derived macrophages out of all leukocytes in the ischemic right hemisphere of sham controls, and mice induced with MCAO who were either injected with PBS, 10M aMOG CD4/CD8 CAR T cells, or 10M aMOG CD4 CAR T cells immediately following MCAO. Statistical significance was determined using an ordinary one-way ANOVA with Dunnett’s multiple comparisons test. ∗∗p≤0.01, ∗∗∗∗p≤0.0001. (K) Boxplot showing frequencies of monocyte-derived macrophages out of all leukocytes in the non-ischemic left hemisphere of sham controls, and mice induced with MCAO who were either injected with PBS, 10M aMOG CD4/CD8 CAR T cells, or 10M aMOG CD4 CAR T cells immediately following MCAO. Statistical significance was determined using an ordinary one-way ANOVA with Dunnett’s multiple comparisons test. ∗∗∗p≤0.001. (L) Boxplot showing frequencies of cDC2 cells out of all leukocytes in the ischemic right hemisphere of sham controls, and mice induced with MCAO who were either injected with PBS, 10M aMOG CD4/CD8 CAR T cells, or 10M aMOG CD4 CAR T cells immediately following MCAO. Statistical significance was determined using an ordinary one-way ANOVA with Dunnett’s multiple comparisons test. ∗∗p≤0.01. (M) Boxplot showing frequencies of cDC2 cells out of all leukocytes in the non-ischemic left hemisphere of sham controls, and mice induced with MCAO who were either injected with PBS, 10M aMOG CD4/CD8 CAR T cells, or 10M aMOG CD4 CAR T cells immediately following MCAO. Statistical significance was determined using an ordinary one-way ANOVA with Dunnett’s multiple comparisons test. ∗∗∗p≤0.001. (N) Boxplot showing frequencies of activated DCs out of all leukocytes in the ischemic right hemisphere of sham controls, and mice induced with MCAO who were either injected with PBS, 10M aMOG CD4/CD8 CAR T cells, or 10M aMOG CD4 CAR T cells immediately following MCAO. Statistical significance was determined using an ordinary one-way ANOVA with Dunnett’s multiple comparisons test. ∗p≤0.05, ∗∗p≤0.01. (O) Boxplot showing frequencies of activated DCs out of all leukocytes in the non-ischemic left hemisphere of sham controls, and mice induced with MCAO who were either injected with PBS, 10M aMOG CD4/CD8 CAR T cells, or 10M aMOG CD4 CAR T cells immediately following MCAO. Statistical significance was determined using an ordinary one-way ANOVA with Dunnett’s multiple comparisons test. ∗p≤0.05.

Intravenous administration of 10 million aMOG CD4/CD8 (mixture of CD4⁺ and CD8⁺) CAR T cells immediately after MCAO resulted in preferential accumulation after 10 days within the damaged right hemisphere, with minimal presence in the spleen and lung and substantially less in the left hemisphere (Figure 2D). Immunohistochemistry confirmed enrichment of T cells within the infarcted region (Figure S2C). We decided to compare CD4/CD8 CAR T versus CD4⁺ CAR T alone. We found that although CD4/CD8 CAR T cells reached similar frequencies as CD4⁺ CAR T cells in the injured hemisphere, as assessed by flow cytometry, CD4⁺ CAR T cells were markedly reduced in the non-ischemic hemisphere (p≤0.01) (Figure 2E), indicating that the CD4⁺ CAR T approach has greater spatial specificity for the damaged site.

To dissect the molecular mechanisms underlying these differences, we performed scRNA-seq of CNS immune cells. CAR T treatment broadly altered the immune landscape relative to both controls and untreated mice that underwent MCAO, increasing infiltration of peripheral immune cells (Figure 2F). However, shifts in specific immune populations occurred in a T cell subset dependent manner (Figure 2F). While CD4⁺ CAR T treatment led to an overall reduction in peripheral leukocyte infiltration, the immune cells that did enter the CNS were largely confined to the right ischemic hemisphere (Figure 2F, S2D), further underscoring the spatial precision with which CD4⁺ CAR T cells target the damaged tissue.

In the microglia compartment, CD4⁺ CAR T treatment reduced overall inflammatory transcriptional signatures relative to CD4/CD8 CAR T cells (Figure 2G). When assessing specific microglia subtypes, CD4⁺ CAR T cells increased the proportion of DAM specifically in the ischemic hemisphere (Figure S2E), while preserving homeostatic microglia in the non-ischemic hemisphere (Figures S2F-H). MHC-II expressing microglia expanded in the injured hemisphere under both CAR T designs (Figure 2H), but were reduced in the non-damaged hemisphere following CD4⁺ CAR T treatment compared with CD4/CD8 CAR T cells (Figure 2I).

Similar hemisphere-specific patterns were observed across myeloid subsets: monocyte-derived macrophages (Figures 2J-K), cDC2s (Figures 2L-M), and activated DCs (Figures 2N-O) all expanded in the ischemic right hemisphere following both CAR T treatments, yet each population was reduced in the non-ischemic hemisphere in CD4⁺ CAR T-treated mice, highlighting the selective localization of CD4⁺ CAR T activity to the site of CNS injury.

Within the lymphocyte compartment, CD4/CD8 CAR T therapy drove strong NK cell recruitment, but CD4⁺ CAR T treatment resulted in markedly lower NK cell infiltration across both hemispheres (Figures S2I-J). Naïve CD4⁺ T cells expanded following both CAR T designs, but most prominently within the injured hemisphere in CD4⁺ CAR T treated mice (Figures S2K-L). Serum analyses revealed no systemic cytokine increases one week following CD4/CD8 CAR T administration (Figure S2M).

Together, these results demonstrate that whereas CD4/CD8 CAR T cells broadly amplify post-ischemic immune infiltration, CD4-restricted CAR T cells achieve a more spatially precise and reparative immune reprogramming of the injured hemisphere, reshaping microglial and myeloid states while minimizing off-target activation.

### BDNF-secreting CAR T cells reprogram the neuroinflammatory microenvironment, enhance Treg responses, and improve functional recovery after stroke

While CD4-restricted CAR T cells achieved spatially precise immune reprogramming of the injured hemisphere, emerging evidence indicates that CD8⁺ T cells can also exert protective and disease-modifying roles in neurodegenerative contexts through direct interactions with microglia and modulation of their functional states^19–21^. Thus, restricting therapeutic design exclusively to CD4⁺ compartments may overlook complementary and beneficial functions mediated by CD8⁺ T cells. Motivated by this hypothesis, we next asked whether we could retain the advantageous precision of CD4 focused approaches while also leveraging the protective potential of CD8⁺ T cells. To achieve this, we engineered cargo releasing aMOG CAR T cells, specifically, CD4⁺ and CD8⁺ CAR T cells secreting the neurotrophic factor BDNF (Figure 3A), with the goal of reshaping the post-ischemic microenvironment toward repair, minimizing cytotoxicity, and harnessing the combined reparative capacities of both T cell subsets. Constitutive BDNF secretion by engineered CAR T cells was first validated *in vitro* (Figure 3B). To confirm that the secreted BDNF retained biological activity, conditioned medium from BDNF-CAR T cells was applied to primary mouse neurons, resulting in robust induction of immediate-early neuronal genes at levels comparable to synthetic BDNF (Figure 3C). BDNF-CAR T cells exhibited activation levels comparable to CAR-only T cells in response to myelin antigen stimulation (Figure S3A), consistent with intact antigen recognition and signaling. This was expected, as BDNF and CAR constructs were introduced via co-transduction rather than through a single P2A-linked construct. Thus, any *in vivo* differences observed following transfer of BDNF-CAR T cells are unlikely to reflect impaired CAR activation and instead can be attributed to the added effects of BDNF secretion.

**Figure 3.**
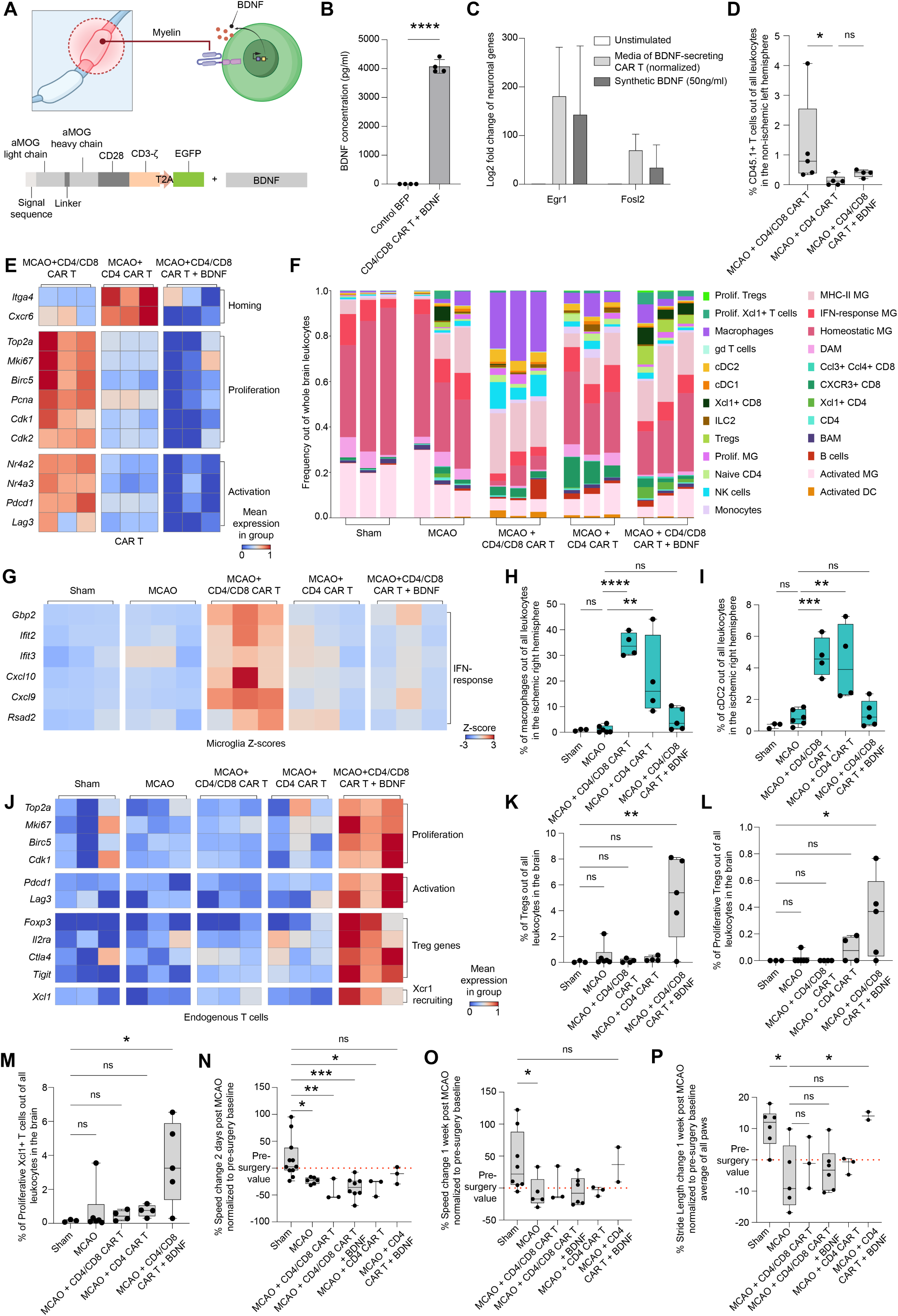
BDNF-secreting CAR T cells reprogram the neuroinflammatory microenvironment, enhance Treg responses, and improve functional recovery after stroke. (A) Schematic illustration of BDNF-secreting aMOG CAR T cells and schematic description of aMOG CAR T and BDNF vectors delivered by co-transduction. (B) Mouse BDNF ELISA assay of BDNF-secreting aMOG CD4/CD8 CAR T cells, and control T cells transduced with BFP. Statistical significance was determined using an unpaired t-test. ∗∗∗∗p≤0.0001. (C) qPCR data of primary mouse neurons that were either unstimulated, conditioned with medium from BDNF-CAR T cells, or stimulated with 50ng/ml synthetic BDNF. (D) Boxplot showing frequency of adoptively transferred CD45.1+ T cells out of all cells in the non-ischemic left hemisphere of mice induced with MCAO who were either treated with 10M aMOG CD4/CD8 CAR T cells, 10M aMOG CD4 CAR T cells, or 10M BDNF-secreting aMOG CD4/CD8 CAR T cells immediately following MCAO. Statistical significance was determined using the Kruskal-Wallis test with Dunn’s multiple comparisons test. ∗p≤0.05. (E) Heatmap showing homing, proliferation, and activation gene expression signatures of CAR T cells in the brains of mice induced with MCAO who were either treated with 10M aMOG CD4/CD8 CAR T cells, 10M aMOG CD4 CAR T cells, or 10M BDNF-secreting aMOG CD4/CD8 CAR T cells immediately following MCAO (n = 9). (F) Bar plots showing the relative abundance of each cell population out of all leukocytes in the brains of sham controls, and mice induced with MCAO who were either injected with PBS, 10M aMOG CD4/CD8 CAR T cells, 10M aMOG CD4 CAR T cells or 10M BDNF-secreting aMOG CD4/CD8 CAR T cells, immediately following MCAO. (G) Heatmap showing IFN-response gene expression of microglia in the brains of sham controls, and mice induced with MCAO who were either injected with PBS, 10M aMOG CD4/CD8 CAR T cells, 10M aMOG CD4 CAR T cells, or 10M BDNF-secreting aMOG CD4/CD8 CAR T cells immediately following MCAO (n = 15). The values are row-normalized z-scores of the pseudobulk gene expression. Each column represents the pooled average reads of a mouse. (H) Boxplot showing frequencies of monocyte-derived macrophages out of all leukocytes in the ischemic right hemisphere of sham controls, and mice induced with MCAO who were either injected with PBS, 10M aMOG CD4/CD8 CAR T cells, 10M aMOG CD4 CAR T cells, or 10M BDNF-secreting aMOG CD4/CD8 CAR T cells immediately following MCAO. Statistical significance was determined using an ordinary one-way ANOVA with Dunnett’s multiple comparisons test. ∗∗p≤0.01, ∗∗∗∗p≤0.0001. (I) Boxplot showing frequencies of cDC2 cells out of all leukocytes in the ischemic right hemisphere of sham controls, and mice induced with MCAO who were either injected with PBS, 10M aMOG CD4/CD8 CAR T cells, 10M aMOG CD4 CAR T cells, or 10M BDNF-secreting aMOG CD4/CD8 CAR T cells immediately following MCAO. Statistical significance was determined using an ordinary one-way ANOVA with Dunnett’s multiple comparisons test. ∗∗p≤0.01, ∗∗∗p≤0.001. (J) Heatmap showing proliferation, activation, Treg genes and Xcr1-recruitment gene expression signatures of endogenous T cells in the brains of sham controls and mice induced with MCAO who were either treated with 10M aMOG CD4/CD8 CAR T cells, 10M aMOG CD4 CAR T cells, or 10M BDNF-secreting aMOG CD4/CD8 CAR T cells immediately following MCAO (n = 15). (K) Boxplot showing frequencies of Tregs out of all leukocytes in the brains of sham controls, and mice induced with MCAO who were either injected with PBS, 10M aMOG CD4/CD8 CAR T cells, 10M aMOG CD4 CAR T cells, or 10M BDNF-secreting aMOG CD4/CD8 CAR T cells immediately following MCAO. Statistical significance was determined using an ordinary one-way ANOVA with Dunnett’s multiple comparisons test. ∗∗p≤0.01. (L) Boxplot showing frequencies of proliferative Tregs out of all leukocytes in the brains of sham controls, and mice induced with MCAO who were either injected with PBS, 10M aMOG CD4/CD8 CAR T cells, 10M aMOG CD4 CAR T cells, or 10M BDNF-secreting aMOG CD4/CD8 CAR T cells immediately following MCAO. Statistical significance was determined using an ordinary one-way ANOVA with Dunnett’s multiple comparisons test. ∗p≤0.05. (M) Boxplot showing frequencies of proliferative Xcl1+ T cells out of all leukocytes in the brains of sham controls, and mice induced with MCAO who were either injected with PBS, 10M aMOG CD4/CD8 CAR T cells, 10M aMOG CD4 CAR T cells, or 10M BDNF-secreting aMOG CD4/CD8 CAR T cells immediately following MCAO. Statistical significance was determined using an ordinary one-way ANOVA with Dunnett’s multiple comparisons test. ∗p≤0.05. (N) Boxplot showing percent speed change 2 days post MCAO surgery, normalized to pre-surgery baselines, and depicting sham controls and mice that underwent MCAO with gait deficits, who were either injected with PBS, 10M aMOG CD4/CD8 CAR T cells, 10M aMOG CD4 CAR T cells, 10M BDNF-secreting aMOG CD4/CD8 CAR T cells or 10M BDNF-secreting aMOG CD4 CAR T cells, immediately following MCAO. Statistical significance was determined using an ordinary one-way ANOVA with Dunnett’s multiple comparisons test. ∗p≤0.05, ∗∗p≤0.01, ∗∗∗p≤0.001. (O) Boxplot showing percent speed change 1 week post MCAO surgery, normalized to pre-surgery baselines, and depicting sham controls and mice that underwent MCAO with gait deficits, who were either injected with PBS, 10M aMOG CD4/CD8 CAR T cells, 10M aMOG CD4 CAR T cells, 10M BDNF-secreting aMOG CD4/CD8 CAR T cells or 10M BDNF-secreting aMOG CD4 CAR T cells, immediately following MCAO. Statistical significance was determined using an ordinary one-way ANOVA with Sidak’s multiple comparisons test. ∗p≤0.05. (P) Boxplot showing percent stride length change 1 week post MCAO surgery, on average across all paws, normalized to pre-surgery baselines, and depicting sham controls and mice that underwent MCAO with gait deficits, who were either injected with PBS, 10M aMOG CD4/CD8 CAR T cells, 10M aMOG CD4 CAR T cells, 10M BDNF-secreting aMOG CD4/CD8 CAR T cells or 10M BDNF-secreting aMOG CD4 CAR T cells, immediately following MCAO. Statistical significance was determined using an ordinary one-way ANOVA with Sidak’s multiple comparisons test. ∗p≤0.05.

We next examined how BDNF secretion influences CAR T cell behavior *in vivo*. BDNF-CAR T cells accumulated in the injured hemisphere at frequencies similar to CD4/CD8 and CD4-restricted CAR T cells (0.4-3.4% out of all leukocytes; Figure S3B), but were markedly reduced in the non-ischemic hemisphere (Figure 3D), mirroring the spatial specificity of CD4⁺ CAR T cells. This suggested that BDNF secretion enhanced site-specific localization to the damaged CNS while minimizing off-target infiltration. To assess how BDNF modulates CAR T phenotype, we profiled CAR T cells in the CNS. BDNF-CAR T cells exhibited reduced *in vivo* proliferation and activation compared to CD4/CD8 CAR T cells, resembling the phenotype of CD4⁺ CAR T cells (Figure 3E). They also displayed an increased naïve-like signature relative to other CAR T designs (Figure S3C). Because CAR T cells or any other immune cell type in our atlas did not express the BDNF receptor TrkB (Figure S3D), which is largely expressed on neurons^54,55^, astrocytes^56^, and oligodendrocytes^57,58^, these phenotypic shifts likely arise from indirect, microenvironment-mediated effects rather than direct BDNF signaling. This indirect effect is consistent with the observation that no differences in proliferation between BDNF-CAR T and other CAR T designs were observed *in vitro* (data not shown).

To dissect the environmental changes driving this reprogramming, we performed scRNA-seq of CNS immune cells following BDNF-CAR T therapy. BDNF-CAR T treatment substantially reshaped the immune landscape compared to both CD4/CD8 and CD4-restricted CAR T therapies (Figures 3F, S3E). In the microglia compartment, BDNF-CAR T cells reduced inflammatory transcriptional signatures similar to CD4⁺ CAR T cells (Figure 3G). Strikingly, BDNF-CAR T treatment increased the abundance of homeostatic microglia in the ischemic hemisphere to levels exceeding those seen with CD4⁺ CAR T cells (Figure S3F).

BDNF-CAR T treatment further attenuated myeloid recruitment to the injury site, resulting in markedly reduced infiltration of macrophages (Figure 3H) and cDC2s (Figure 3I) into the ischemic hemisphere relative to both other CAR T therapies. In the non-ischemic hemisphere, macrophage and cDC2 frequencies remained low, resembling those seen with CD4⁺ CAR T therapy (Figures S3G-H), indicating that BDNF release promotes spatially restricted myeloid reprogramming. Within the T cell compartment, BDNF elicited a pronounced remodeling of endogenous T cell states. BDNF-CAR T treatment increased proliferation, activation, and Treg signatures across endogenous T cells (Figure 3J). Notably, BDNF-CAR T therapy uniquely expanded both Tregs and proliferative Tregs in the CNS, an effect that was not observed with any other CAR T design (Figures 3K-L).

BDNF-CAR T treatment also increased the frequencies of endogenous Xcl1⁺ CD4 and Xcl1⁺ CD8 T cells (Figures S3I-J). Consistent with this, endogenous proliferative Xcl1⁺ T cells were overall more abundant in BDNF-treated mice (Figure 3M). Because the canonical Xcr1⁺ cDC1 population was not expanded (Figures S3K, S1B), these effects are unlikely to be mediated through cDC1s. Instead, they may reflect engagement of Xcr1-expressing cells in the CNS, including neurons, which were shown to upregulate Xcr1 following brain injury^59,60^. Notably, Xcl1 has been shown to enhance neuronal excitability, and Xcl1-Xcr1 signaling has been implicated in brain injury responses^59,60^, suggesting a potential neuro-immune modulatory role for this pathway. Histological assessment showed that the infarct remained clearly detectable after BDNF-CAR T treatment (Figure S3L). Thus, the most striking effects of the therapy were not anatomical but reflected a robust reprogramming of the cellular and molecular landscape within the injured hemisphere.

Next, we combined our two approaches of CD4⁺ CAR T and BDNF secretion, and compared it to CD4⁺ CAR T alone. BDNF-CD4⁺ CAR T cells exhibited reduced *in vivo* proliferation compared to CD4⁺ CAR T cells (Figure S4A). Behaviorally, BDNF-CD4⁺ CAR T treated mice showed superior gait outcomes observed as early as day 2 and sustained at day 7, and were the only group to exceed their pre-surgery performance levels within a week. These mice exhibited increased running speed (Figures 3N-O, S4B) and increased stride length and swing speed across all paws (Figures 3P, S4C-D), approaching values observed in control animals. Further studies will be required to establish the superiority of BDNF-CD4⁺ CAR T treatment across additional models of CNS injury and neurodegeneration.

## Discussion

CAR T cell therapy is increasingly recognized as a transformative modality for neurological diseases^1–5^. Unlike brain tumors or CNS autoimmune diseases, where target-cell elimination is the primary goal, the engineering strategies needed to safely and effectively deploy CAR T cells for tissue repair in neurodegeneration or CNS injury remain largely undefined. Growing evidence shows that endogenous T cells can exert protective functions in neurodegeneration and CNS injury^10–17^. In spinal cord injury, clonally expanded CD4⁺ T cells recognizing myelin and neuronal self-antigens have been shown to promote neuroprotection by modulating myeloid cell states^10^, demonstrating that T cells can support repair rather than exacerbate damage^10,13,14,17^. These findings position CD4⁺ T cells as attractive candidates for CAR T applications in the injured brain. However, emerging work suggests that CD8⁺ T cells can also contribute to neuroprotection through interactions with microglia^19–21^, highlighting the need to systematically evaluate both CD4⁺ and CD8⁺ CAR T programs to fully leverage T cell mediated immune modulation in the CNS. Here, we used ischemic stroke as a proof of concept and designed MOG-targeting CAR T cells, building on prior work showing that myelin-reactive T cells can be neuroprotective in CNS injury^10,13,14,61^. We delineated how distinct CAR architectures, including CD4⁺ and CD8⁺ CAR T cells and CAR T cells engineered to secrete the neurotrophic factor BDNF, differentially shape the post-injury immune microenvironment.

Our findings demonstrated that CAR T cells containing a combination of CD8⁺ and CD4⁺ cells, though capable of trafficking to areas of CNS damage, triggered broad infiltration of peripheral immune cells and widespread activation of microglial and myeloid populations. In contrast, CD4⁺ CAR T cells exhibited markedly higher spatial precision, accumulating preferentially in the injured hemisphere and minimizing off-target immune remodeling. This hemisphere-specific targeting resulted in microglial reprogramming that preserved homeostatic states in the non-damaged hemisphere while promoting DAM and inflammatory resolution specifically within the damaged site.

CAR T cells that proliferate extensively within the CNS risk exacerbating inflammation or enhancing cytotoxic damage^3,4^. Our scRNA-seq profiling revealed that CD4/CD8 CAR T cells adopt a highly activated, proliferative phenotype within the CNS, whereas CD4⁺ CAR T cells maintain a more restrained activation state. BDNF secretion shifted CAR T cells toward an inflammation-resolving phenotype with reduced proliferation, an effect likely tuned through microenvironmental modulation. BDNF delivery reshaped microglial and myeloid states and reduced macrophage and dendritic cell infiltration into the injured hemisphere, compared to CD4/CD8 CAR T treatment. While BDNF-CAR T cells resembled the spatial specificity of CD4⁺ CAR T cells, they also induced unique lymphocyte programs, including enhanced recruitment and proliferation of endogenous Tregs and increased Xcl1⁺ CD4 and CD8 T cells, populations potentially acting through Xcl1/Xcr1 pathways implicated in neuronal excitability and brain injury responses^59,60^.

Phenotypically, CD4⁺ BDNF-CAR T therapy significantly improved gait recovery, achieved through cellular and molecular reprogramming. While our study implicates neuroimmune circuits underlying functional recovery, future studies will explore the functional interplay between engineered CAR T cells and neuronal survival and plasticity, as the primary cells expressing the BDNF receptor TrkB are not immune cells, but rather neurons^54,55^, astrocytes^56^, and oligodendrocytes^57,58^. Collectively, our findings highlight that the therapeutic potential of CAR T cells in neurodegeneration and CNS injury lies in their ability to home precisely to sites of damage, engage with and reprogram local immune and neural cells, and serve as modular platforms for delivering reparative factors. These capabilities position CAR T therapy as a versatile tool for neurological disease, one that can be engineered not only to eliminate pathogenic cells, but to sculpt a microenvironment conducive to repair. Our study provides a roadmap for CNS targeting CAR T strategies and supports the continued development of CNS-tailored, cargo-delivering CAR T cells as a new class of reparative therapeutics.

## Resource availability

### Lead contact

Further information and requests for resources and reagents should be directed to and will be fulfilled by the lead contact, Ido Amit.

### Materials availability

Correspondence and requests for materials should be addressed to Ido Amit. All biological materials can be obtained from the corresponding authors following reasonable requests.

### Data and code availability

Single-cell RNA-seq data will be deposited in the Gene Expression Omnibus and will be made publicly available as of the date of publication.

Any additional information required to re-analyze the data reported in this paper is available from the lead contact upon request.

## Acknowledgments

We would like to thank Ms. Tal Bigdary for her contribution to the visual presentation. I.A. is an Eden and Steven Romick Professorial Chair, supported by the HHMI International Scholar Award, funded by the European Union ERC advanced grant (no. 101055341-TROJAN-Cell), European Union (IMMEDIATE EU#: 101095540), the MBZUAI/WIS joint program on artificial intelligence. Lotte and John Hecht Memorial Foundation and the Schwartz Reisman Collaborative Science Program, the Deutsche Forschungsgemeinschaft (DFG, German Research Foundation) – Project-ID 259373024 – TRR 167, the Israel Science Foundation grant no. 1944/22, the European Union (ERC, MiTE, 101123436). This research was supported by a Research Professorship grant from the Israel Cancer Research Fund, United States-Israel Binational Science Foundation (BSF), Jerusalem, Israel, Ministry of Innovation, Science & Technology (1001703362), Dwek Institute for Cancer Therapy Research, Moross Integrated Cancer Center, EKARD Institute for Cancer Diagnosis Research, Morris Kahn Institute for Human Immunology, Swiss Society Institute for Cancer Prevention Research, Elsie and Marvin Dekelboum Family Foundation, an ASPIRE Award from The Mark Foundation for Cancer Research, Alzheimer’s Association (ABA-25-1373817). This study was supported by the Center for Immunotherapy at the Weizmann Institute of Science. This work was supported by the AI Hub at the Institute for Artificial Intelligence, Weizmann Institute of Science. R.S. was supported by NeuroMac School (CRC/TRR167) and Azrieli Institute for Brain and Neural Sciences.

## Author contributions

R.S. calibrated and developed experimental protocols; conceptualized, designed, performed, and analyzed experiments and interpreted results; performed all bioinformatic analyses; wrote the manuscript; and directed the project. M.B.Y. calibrated and developed experimental protocols; conceptualized, designed, performed, and analyzed experiments and interpreted results; and wrote the manuscript. P.B. consulted and wrote the manuscript. Y.F. performed experiments. Y.K. performed the MCAO and sham surgeries. M.T. consulted on the gait motor assessment. V.K. developed and performed the TOVI imaging. O.B. performed the histological analyses. E.D. and K.M. performed the scRNA-seq library preparation, sequencing, and read alignment. R.M. interpreted results and wrote the manuscript. J.K. conceptualized, designed, interpreted results and wrote the manuscript. I.A. developed experimental protocols, mentored and directed the project, conceptualized, and designed experiments, interpreted results, and wrote the manuscript.

## Declaration of generative AI and AI-assisted technologies in the writing process

During the preparation of this work, the authors used ChatGPT in order to improve writing clarity. After using this tool, the authors reviewed and edited the content as needed and take full responsibility for the content of the publication.

## Materials and Methods

### Retroviral vector production

Platinum-E (Plat-E) cells were cultured in 15cm tissue culture plates in Dulbecco’s Modified Eagle’s Medium (DMEM) supplemented with 10% heat-inactivated fetal bovine serum (FBS), 1% sodium pyruvate, penicillin-streptomycin, and L-glutamine (referred to as complete DMEM). Cells were maintained at 60–70% confluency. One day prior to transfection, Plat E cells were seeded at 1×10⁶ cells per well into 6-well plates and incubated overnight. On the day of transfection, when cells reached over 60–70% confluency, the medium was replaced with DMEM without antibiotics. For each well, cells were transfected using a mixture prepared from 10 µL of Lipofectamine 2000 (Cat. No. 11668027, Thermo Fisher) that was diluted in 240 µL of Opti-MEM medium. Separately, 1 µg of the retroviral vector of interest and 0.5 µg of the packaging plasmid pCL-Eco (Addgene, Cat. No. 12371) were combined in Opti-MEM to a total volume of 250 µL. After 5 minutes of separate incubation, the Lipofectamine mixture was gently combined with the plasmid mixture by inverting the tube and incubated at room temperature for 20 minutes. The entire 500 µL transfection mixture was then added dropwise to each well of Plat E cells. Cells were incubated at 37°C for 5 to 8 hours, after which the medium was replaced with fresh complete DMEM. Viral supernatants were collected 48 hours post-transfection and filtered through a 0.45 µm syringe filter to remove cellular debris.

### T lymphocyte isolation and activation

Mouse spleens were harvested and mechanically dissociated to generate single-cell suspensions, and RBC were lysed with ACK buffer (Gibco, Cat. No. A1049201). Total T lymphocytes or CD4 T cells were subsequently isolated from the total splenocyte population using the magnetic bead separation kits - Pan T Cell Isolation Kit (Miltenyi Biotec, Cat. No. 130-095-130) for all T cells or CD4+ T Cell Isolation Kit (Miltenyi Biotec, Cat. No. 130-095-248) for CD4 T cells, according to the manufacturer’s instructions. Isolated T cells were cultured in RPMI-1640 medium supplemented with 10% heat-inactivated fetal bovine serum (FBS), 1% sodium pyruvate, penicillin-streptomycin, L-glutamine, and non-essential amino acids, 0.1% β-mercaptoethanol, referred to as complete RPMI medium. 10 ng/mL recombinant murine IL-2 (PeproTech, Cat. No. 212-12-100), IL-7 (PeproTech, Cat. No. 217-17-100), and IL-15 (PeproTech, Cat. No. 210-15-100) have been added to the medium, hereafter referred to as lymphocyte medium. For T cell activation, 24-well tissue culture-treated plate was coated overnight at 4°C (or for 2 hours at 37°C) with 250 µL per well of anti-CD3 antibody (Bio Legend, Cat. No. 100340) diluted to 1 ug/mL in sterile phosphate-buffered saline (PBS). Purified T cells were then seeded into the coated activation plates at a density of 3×10⁶ cells per well in lymphocyte medium containing soluble anti-CD28 antibody (Bio Legend, Cat. No. 102116) at 2 ug/mL. Plates were incubated at 37°C for 24 hours to induce activation.

### Retroviral transduction

For retroviral transduction, 24-well non-tissue culture-treated plates were coated with 25 µg/mL RetroNectin (Takara, Cat. No. T100A) in PBS at 0.5 mL per well. Coating was performed for 2 hours at room temperature or overnight at 4°C. Following coating, plates were blocked with PBS containing 2% bovine serum albumin (BSA) for 30 minutes at room temperature, then washed once with Hanks’ Balanced Salt Solution (HBSS) supplemented with 2.5% 1M HEPES. One milliliter of each viral supernatant type (for single or double transductions) was added gently to the coated wells and centrifuged at 2,000 × g for 2 hours at 32°C to facilitate virus adsorption onto the RetroNectin-coated surface. After virus adsorption, viral supernatants were carefully removed. Activated T cells (1×10⁶ cells per well) in 1 mL of lymphocyte medium were then gently added to each well by pipetting along the wall to avoid disturbing the RetroNectin coating. Plates were centrifuged at 350 × g for 20 minutes at 32°C to promote cell adherence and incubated at 37°C. Cells were subsequently collected and transferred to new tissue culture-treated plates at a density of 1×10⁶ cells/ml every 48 hours until further use.

### IFNG and BDNF ELISA

To quantify Ifng levels secreted by CAR T cells in response to myelin stimulation, transduced cells were collected and counted. Cells were then centrifuged at 350 × g for 5 minutes at 4°C. Following centrifugation, cells were seeded into 96-well tissue culture-treated plates for 24 hours in complete RPMI medium, deprived of cytokines with or without myelin stimulation. After the 24-hour incubation period, cell supernatants were collected. Ifng secretion levels were measured using a mouse Ifng ELISA kit (BioLegend ELISA MAX™ Deluxe Set Mouse IFNg, Cat. No. 430816) according to the manufacturer’s protocol. Absorbance at 450 nm was immediately recorded using a spectrophotometer. Background absorbance, determined from negative control samples, was subtracted from all readings. Absolute Ifng levels were calculated by interpolation from a standard curve generated using the Ifng standards provided in the ELISA kit. BDNF ELISA assay was performed to assess secretion by CAR T cells (by R&D Systems Human/Mouse BDNF DuoSet ELISA, Cat. No. DY248), according to the manufacturer’s protocol. Absorbance measurement and analysis were done as described for the Ifng ELISA.

### Tissue processing

Mice were transcardially perfused with sterile PBS, and brains, spleens, and lungs were extracted. Brains were immediately placed into 1 ml volume of respective digestion buffers. Brain digestion buffer contained 100 µg/mL DNase I (Sigma-Aldrich, Cat. No. 10104159001) and 200 µg/mL Collagenase IV (Enco, Cat. No. LS004188) in PBS. Tissues were mechanically minced in their respective buffers. Brains were then incubated with shaking at 37°C for 30 minutes. During the incubation period, samples were further mechanically dissociated by trituration through a 21G needle. Digested tissue mixes were filtered through a 40 µm cell strainer and washed with PBS to inhibit enzyme activity. Brain cell suspensions were centrifuged at 350 × g for 5 minutes at 4°C. To isolate mononuclear cells from CNS tissues, cell pellets were resuspended in 8 mL of a Percoll solution (30% Percoll, 60% double-distilled water (DDW), and 10% 10x PBS). Samples were centrifuged at 2000 rpm for 20 minutes at 4°C, with an acceleration speed of 4 and a brake speed of 3. After careful removal of the supernatant and myelin layer, the remaining cell pellets were washed in MACS buffer (PBS with 0.5% BSA and 2 mM EDTA). Lungs were harvested and mechanically minced in 5 mL of pre-heated (37°C) digestion buffer consisting 0.0125 mg/mL DNase I and 1 mg/mL Collagenase IV in plain RPMI-1640 medium. Tissues underwent two rounds of mechanical dissociation using a gentleMACS dissociator (Miltenyi Biotec, Cat. No. 130-093-235), with an 8-minute incubation in a 37°C water bath after each round. Following digestion, cell suspensions were passed through a 70µm cell strainer and washed with ice-cold PBS, and red blood cells were lysed with ACK buffer (Gibco, Cat. No. A1049201). Spleens were harvested and mechanically dissociated to generate single-cell suspensions, and red blood cells were lysed with ACK buffer (Gibco, Cat. No. A1049201).

### Flow cytometry staining, analysis, and sorting

Isolated single-cell suspensions from MCAO tissues were stained for surface markers. Cells were incubated in 100µL of MACS buffer with the following antibodies at a 1:100 dilution: anti-CD45.1 PE-Cy7 (clone A20, Cat. No. 110730, BioLegend), anti-CD45.2 APC (clone 104, Cat. No. 558702, BD Biosciences), and anti-TCRb APC-Cy7 (clone H57-597, Cat. No. 109219, BioLegend). All stainings were conducted for 30 minutes on ice in the dark. After staining, cells were washed with cold MACS buffer and filtered through a 70µm cell strainer. To detect cell nuclei and assess viability, 4′,6-diamidino-2-phenylindole (DAPI) was added to all samples 2 minutes before commencing analysis or sorting. Cells were aquired on a BD FACS Symphony 6 flow cytometer (BD Biosciences, San Jose, CA). Flow cytometry data analysis was performed using BD FACSDIVA (BD Biosciences) and the FlowJo (FlowJo LLC version 10) software. Single cells were sorted (total CD45.1, total CD45.2 and CAR T GFP cell enrichment) into 384-well capture plates. Each well contained lysis buffer, 3µL of mineral oil, and 20nM barcoded poly (T) reverse transcription primers. Following sorting, plates were centrifuged to ensure proper cell contact with the lysis solution, then immediately frozen on dry ice and stored at −80°C until further processing.

### Single-cell RNA sequencing (scRNA-seq) library preparation, sequencing, and read alignment

Single-cell RNA sequencing libraries were generated using SPID-seq^62^, a modified version of the MARS-seq protocol^63^. Briefly, individual cells were sorted into 384-well plates as described above. Polyadenylated mRNA from single cells was captured and barcoded during reverse transcription into cDNA. The resulting cDNA was then pooled, fragmented, and amplified to produce sequencing-ready libraries. Quality control was performed for each plate-derived library. Pooled libraries were sequenced on an Illumina NovaSeq X Plus platform at a depth of 10,000-50,000 reads per cell. Reads with identical unique molecular identifiers (UMIs) were collapsed to represent individual RNA molecules. Batch quality was validated by confirming low cross-cell contamination (<3%), assessed by the frequency of spurious UMIs in empty wells. Read alignment was performed using the MARS-seq 2.0 pipeline. Low-quality reads were removed, and the remaining reads were mapped to the mouse reference genome mm10 or to the human reference genome hg38 using HISAT (version 0.1.6), excluding reads with multiple mapping positions. Only exonic reads were counted, based on UCSC gene annotations. UMI uniqueness was confirmed within a 3 kb window. For overlapping exons from distinct genes on the same strand, reads were assigned to a merged gene ID combining both gene symbols.

### Plasmid design and synthesis

To generate the aMOG CAR, the heavy and light chains of the antigen recognition domains of aMOG303 antibody were utilized as binding domains^53^. The aMOG CAR construct was constructed by assembling a single-chain variable fragment (scFv) in a VL-linker-VH orientation, followed by the CD28 transmembrane domain, and the CD3ζ cytoplasmic domain. The CAR sequence was fused to a T2A-EGFP reporter for visualization and tracking. The CAR and BDNF cargo were cloned into the MSGV-1D3-28Z All ITAMs intact retroviral backbone (Addgene, Cat. No. 107226)^64^. The complete plasmid sequences were ordered as double-stranded genes from GenScript.

### Transcranial optical vascular imaging (TOVI)

All animal procedures were approved by the Institutional Animal Care and Use Committee of the Weizmann Institute and were conducted in accordance with institutional and national guidelines for the humane care and use of laboratory animals. Mice were anesthetized by intraperitoneal injection of ketamine (10 mg/kg) and xylazine (100 mg/kg) diluted in PBS to a final volume of 200 µl. A midline incision was made, and the skin over the frontal, temporal, occipital, and parietal regions was removed by blunt dissection. The exposed skull was kept constantly moist with sterile saline. Animals were positioned on a custom-designed stereotaxic holder under the microscope objective. Transcranial optical vascular imaging (TOVI) and image analysis were performed according to the protocol described by Kalchenko et al. (2014)^49^. The duration of the procedure was less than 1.5 hours, after which mice were euthanized by barbiturate overdose.

### Middle cerebral artery occlusion (MCAO)

Focal cerebral ischemia was induced by transient middle cerebral artery occlusion using the intraluminal filament technique. Mice were anesthetized with ketamine and xylazine as described above, the fur over the ventral neck was shaved, and the skin was disinfected. Body temperature was maintained at 37°C throughout surgery and occlusion using a controlled heating pad. A midline cervical incision was made, and the soft tissues were bluntly dissected to expose the carotid arteries. MCAO was performed as described by Engel et al. (2011)^48^. Briefly, a silicone-coated monofilament (702034PK5Re, Doccol Corporation, Sharon, MA, USA; tip diameter 0.20 mm) was introduced into the left internal carotid artery and advanced to occlude the origin of the left middle cerebral artery at the circle of Willis. Occlusion was maintained for 60 minutes while animals were kept on a heated surface, and the surgical field was kept moist and covered with sterile gauze. Additional anesthesia was administered if required. After 60 minutes, the filament was withdrawn to allow reperfusion, and the vessels and skin were sutured. Mice were then transferred to temperature-controlled (37°C) recovery cages and monitored daily, including body weight assessment and administration of analgesics and 1ml saline subcutaneously for 3 days. Sham-operated animals underwent the same surgical procedure without insertion of the occluding filament.

### CatWalk gait analysis

Gait performance was assessed using the CatWalk XT automated gait analysis system (Noldus Information Technology) following MCAO. The system consists of a glass walkway positioned above a high-resolution camera; light is internally reflected within the glass and is emitted only at points of paw contact, resulting in illuminated paw prints that are captured as video frames. As rodents traverse the walkway, the CatWalk XT software quantifies paw print intensity and spatial-temporal parameters of gait. The testing protocol followed Garrick et al. (2021)^51^ with minor modifications. Each mouse was habituated to the apparatus, and two baseline recordings were acquired on separate days before MCAO to characterize normal, baseline gait. After surgery, gait was recorded on days 2, 4, and 7 post-occlusion. For each session, animals were allowed to cross the walkway until at least three compliant runs (uninterrupted crossing at a consistent speed) were obtained. From the CatWalk XT output, the following outcome measures relevant to MCAO-induced motor deficits were extracted and analyzed: run duration, average walking speed, swing speed, and stride length.

### Quantification and statistical analysis

Data analyses were performed as indicated in the relevant Methods sections or by using GraphPad Prism (GraphPad Software).

For normal distributions, one-way analysis of variance (ANOVA) with Dunnett’s multiple comparisons test was used to compare three or more groups, or ordinary two-way ANOVA with Dunnett’s multiple comparisons test was used for two distinct factors (paw and treatment group). When two groups were compared, an unpaired or paired two-tailed student’s t-tests were used to determine statistical significance. When data were not normally distributed, a nonparametric two-tailed Mann-Whitney U test was used when two groups were compared.

Statistical significance indicated in all figures is as follows: ∗p≤0.05, ∗∗p≤0.01, ∗∗∗p≤0.001, ∗∗∗∗p≤0.0001, ns: non-significant.

---

**Figure S1.**
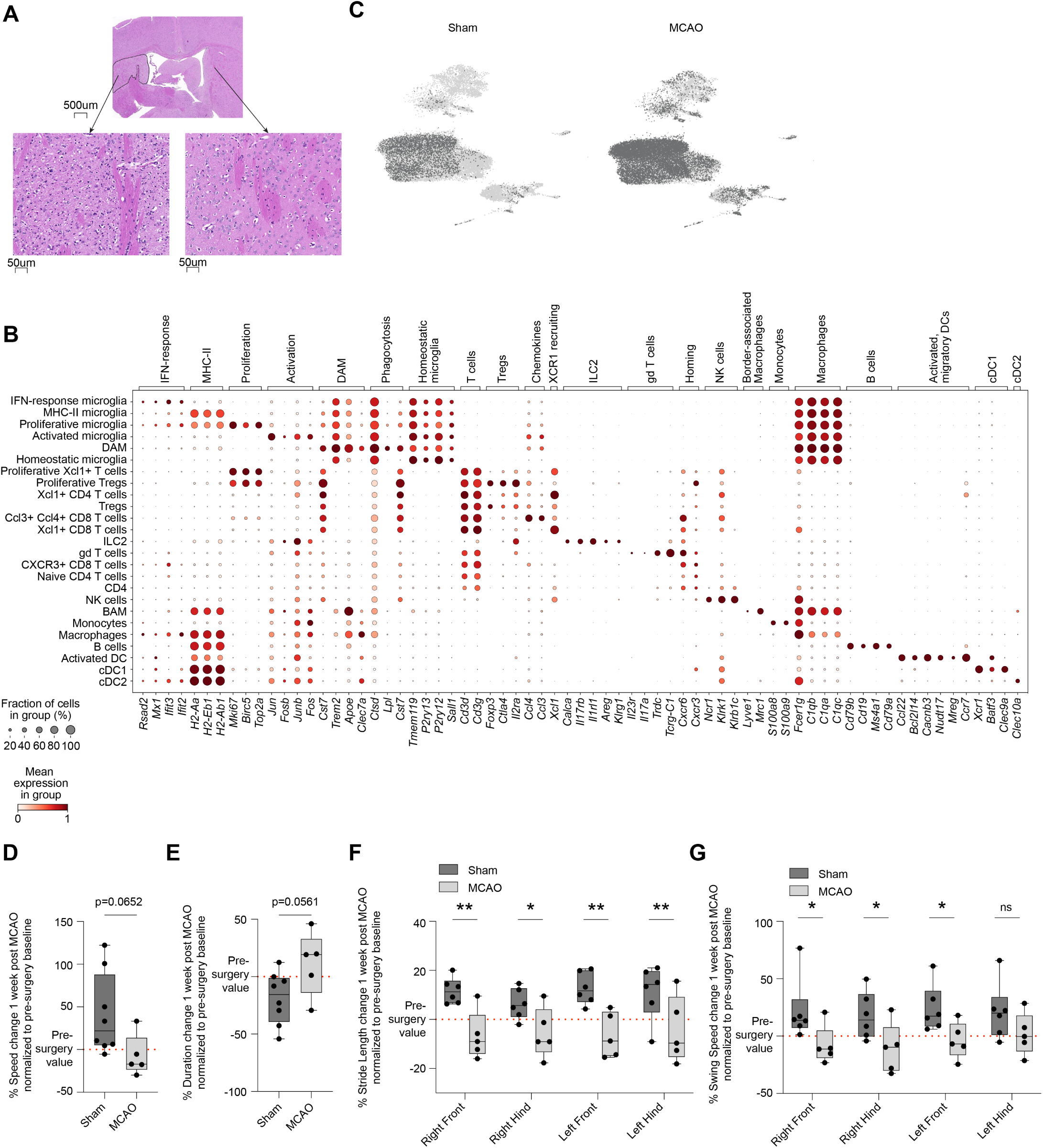
Characterization of the MCAO model, related to Figure 1. (A) H&E staining of a representative mouse brain 8 days post 60 minute occlusion of the MCA. Unilateral moderate vacuolation and scattered red neurons are seen. (B) Dot plot showing normalized gene expression profiles of selected marker genes across cell types in the brain of mice induced with MCAO. Dot size represents the percentage of cells expressing the gene, and color represents the scaled mean gene expression of the cluster. (C) UMAP visualization of scVI embedding on single cells obtained from the brain and integrated across mice induced with MCAO (right) and sham controls (left). (D) Boxplot showing percent speed change 1 week post MCAO surgery, normalized to pre-surgery baselines, and depicting mice that underwent MCAO with gait deficits and controls. Statistical significance was determined using an unpaired t-test. (E) Boxplot showing percent duration change 1 week post MCAO surgery, normalized to pre-surgery baselines, and depicting mice that underwent MCAO with gait deficits and controls. Statistical significance was determined using an unpaired t-test. (F) Boxplot showing percent stride length change 1 week post MCAO surgery, normalized to pre-surgery baselines, and depicting mice that underwent MCAO with gait deficits and controls. Statistical significance was determined using an ordinary two-way ANOVA with Sidak’s multiple comparisons test. ∗∗p≤0.01, ∗p≤0.05. (G) Boxplot showing percent swing speed change 1 week post MCAO surgery, normalized to pre-surgery baselines, and depicting mice that underwent MCAO with gait deficits and controls. Statistical significance was determined using an ordinary two-way ANOVA with Sidak’s multiple comparisons test. ∗p≤0.05.

**Figure S2.**
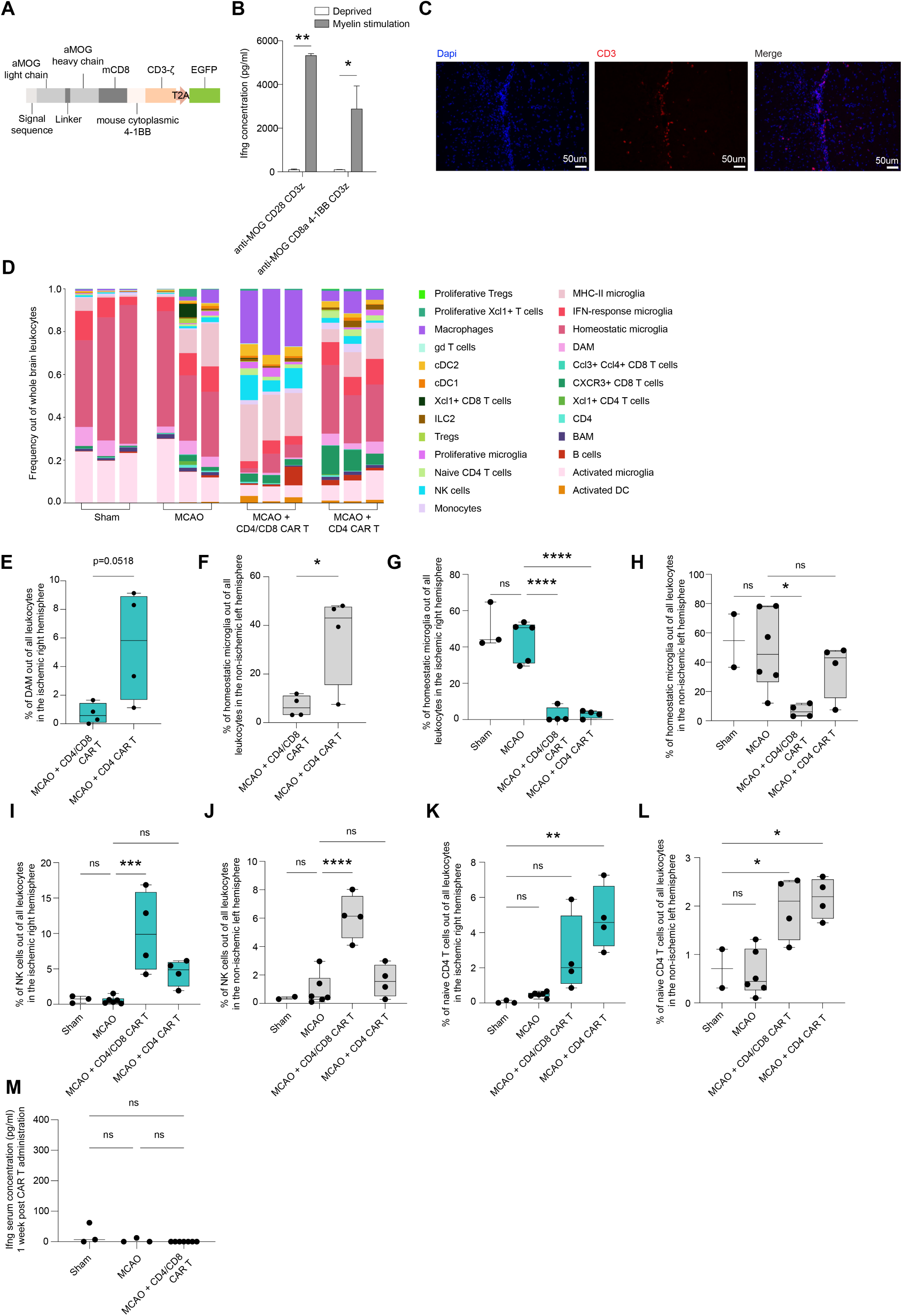
CAR T cells reprogram the post-ischemic CNS, related to Figure 2. (A) Schematic description of aMOG CD8a 4-1BB CD3z CAR vector. (B) Mouse IFN-γ ELISA assay comparing two aMOG CAR variants, either unstimulated or stimulated with myelin. Statistical significance was determined using an ordinary one-way ANOVA with Tukey’s multiple comparisons test. ∗p≤0.05, ∗∗p≤0.01. (C) Immunohistochemistry of a representative section in the right ischemic hemisphere, following an H&E section where an infarct was identified. Staining for CD3 (red) and dapi (blue). (D) Bar plots showing the relative abundance of each cell population out of all leukocytes in the brains of sham controls, and mice induced with MCAO who were either injected with PBS, 10M aMOG CD4/CD8 CAR T cells, or 10M aMOG CD4 CAR T cells immediately following MCAO. (E) Boxplot showing frequencies of DAM cells out of all leukocytes in the ischemic right hemisphere, for mice that underwent MCAO and were treated with either 10M CD4/CD8 CAR T or 10M CD4 CAR T. Statistical significance was determined using an unpaired t-test. (F) Boxplot showing frequencies of homeostatic microglia out of all leukocytes in the non-ischemic left hemisphere, for mice that underwent MCAO and were treated with either 10M CD4/CD8 CAR T or 10M CD4 CAR T. Statistical significance was determined using an unpaired t-test. ∗p≤0.05. (G) Boxplot showing frequencies of homeostatic microglia out of all leukocytes in the ischemic right hemisphere of sham controls, and mice induced with MCAO who were either injected with PBS, 10M aMOG CD4/CD8 CAR T cells, or 10M aMOG CD4 CAR T cells immediately following MCAO. Statistical significance was determined using an ordinary one-way ANOVA with Dunnett’s multiple comparisons test. ∗∗∗∗p≤0.0001. (H) Boxplot showing frequencies of homeostatic microglia out of all leukocytes in the non-ischemic left hemisphere of sham controls, and mice induced with MCAO who were either injected with PBS, 10M aMOG CD4/CD8 CAR T cells, or 10M aMOG CD4 CAR T cells immediately following MCAO. Statistical significance was determined using an ordinary one-way ANOVA with Dunnett’s multiple comparisons test. ∗p≤0.05. (I) Boxplot showing frequencies of NK cells out of all leukocytes in the ischemic right hemisphere of sham controls, and mice induced with MCAO who were either injected with PBS, 10M aMOG CD4/CD8 CAR T cells, or 10M aMOG CD4 CAR T cells immediately following MCAO. Statistical significance was determined using an ordinary one-way ANOVA with Dunnett’s multiple comparisons test. ∗∗∗p≤0.001. (J) Boxplot showing frequencies of NK cells out of all leukocytes in the non-ischemic left hemisphere of sham controls, and mice induced with MCAO who were either injected with PBS, 10M aMOG CD4/CD8 CAR T cells, or 10M aMOG CD4 CAR T cells immediately following MCAO. Statistical significance was determined using an ordinary one-way ANOVA with Dunnett’s multiple comparisons test. ∗∗∗∗p≤0.0001. (K) Boxplot showing frequencies of naive CD4 T cells out of all leukocytes in the ischemic right hemisphere of sham controls, and mice induced with MCAO who were either injected with PBS, 10M aMOG CD4/CD8 CAR T cells, or 10M aMOG CD4 CAR T cells immediately following MCAO. Statistical significance was determined using an ordinary one-way ANOVA with Dunnett’s multiple comparisons test. ∗∗p≤0.01. (L) Boxplot showing frequencies of naive CD4 T cells out of all leukocytes in the non-ischemic left hemisphere of sham controls, and mice induced with MCAO who were either injected with PBS, 10M aMOG CD4/CD8 CAR T cells, or 10M aMOG CD4 CAR T cells immediately following MCAO. Statistical significance was determined using an ordinary one-way ANOVA with Dunnett’s multiple comparisons test. ∗p≤0.05. (M) Mouse IFN-γ ELISA assay of serum of sham controls and mice induced with MCAO who were either injected with PBS or 10M aMOG CD4/CD8 CAR T cells, 1 week following CAR T administration.

**Figure S3.**
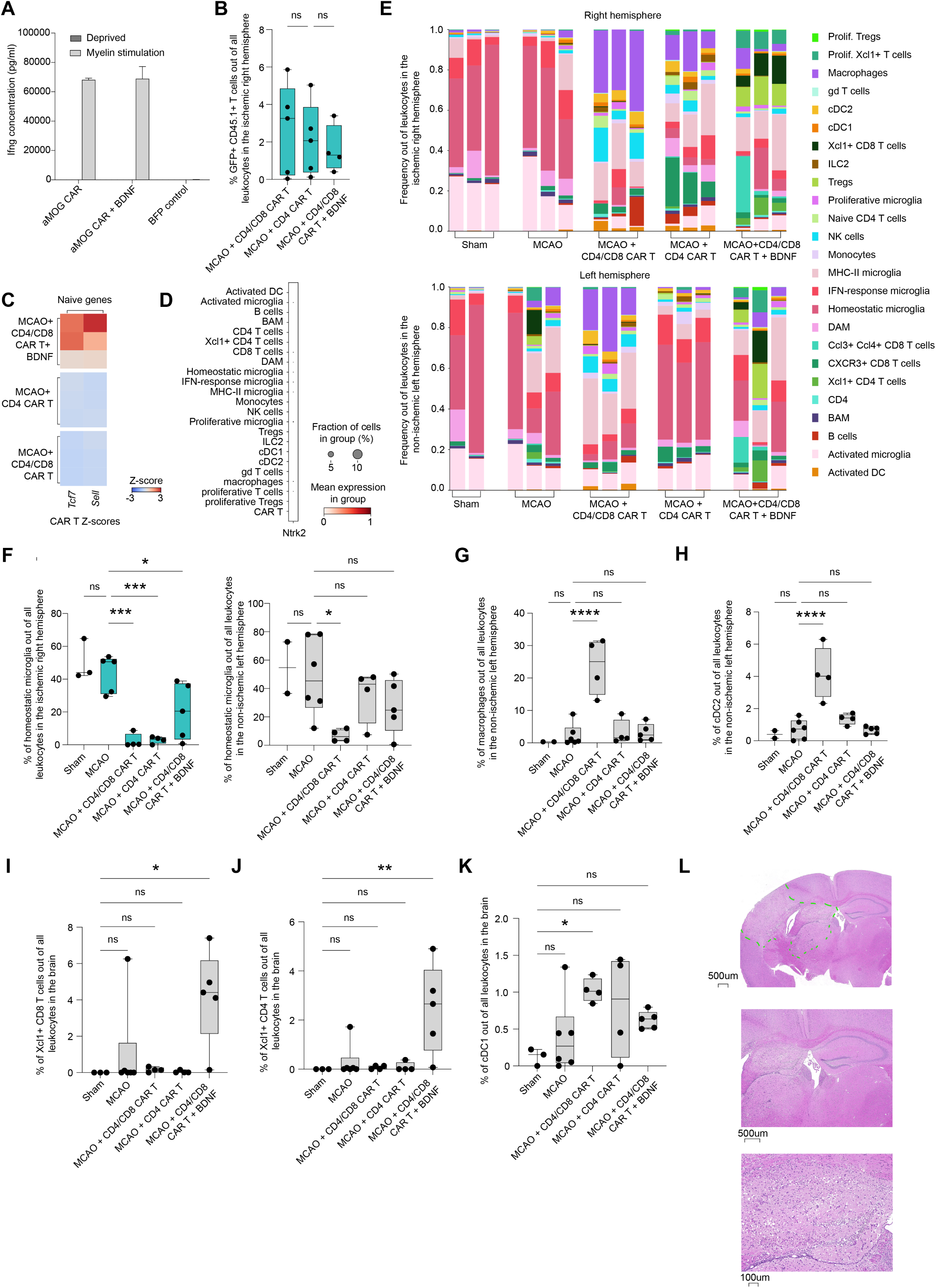
BDNF-secreting CAR T cells reprogram the post-ischemic CNS, related to Figure 3. (A) Mouse IFN-γ ELISA assay comparing aMOG CAR and BDNF-secreting aMOG CAR, either unstimulated or stimulated with myelin. (B) Boxplot showing frequency of adoptively transferred GFP+ CD45.1+ T cells out of all cells in the ischemic right hemisphere of mice induced with MCAO who were either treated with 10M aMOG CD4/CD8 CAR T cells, 10M aMOG CD4 CAR T cells, or 10M BDNF-secreting aMOG CD4/CD8 CAR T cells immediately following MCAO. Statistical significance was determined using an ordinary one-way ANOVA with Dunnett’s multiple comparisons test. (C) Heatmap showing naive gene expression signatures of CAR T cells in the brains of mice induced with MCAO who were either treated with 10M aMOG CD4/CD8 CAR T cells, 10M aMOG CD4 CAR T cells, or 10M BDNF-secreting aMOG CD4/CD8 CAR T cells immediately following MCAO (n = 9). (D) Dot plot showing normalized *Ntrk2* (TrkB) gene expression profiles of cell types in the brains of mice induced with MCAO. Dot size represents the percentage of cells expressing the gene, and color represents the scaled mean gene expression of the cluster. (E) Bar plots showing the relative abundance of each cell population out of all leukocytes in the ischemic right hemisphere (top) and non-ischemic left hemisphere (bottom) of sham controls, and mice induced with MCAO who were either injected with PBS, 10M aMOG CD4/CD8 CAR T cells, 10M aMOG CD4 CAR T cells or 10M BDNF-secreting aMOG CD4/CD8 CAR T cells, immediately following MCAO. (F) Boxplots showing frequencies of homeostatic microglia out of all leukocytes in the ischemic right and non-ischemic left hemispheres of sham controls, and mice induced with MCAO who were either injected with PBS, 10M aMOG CD4/CD8 CAR T cells, 10M aMOG CD4 CAR T cells, or 10M BDNF-secreting aMOG CD4/CD8 CAR T cells immediately following MCAO. Statistical significance was determined using an ordinary one-way ANOVA with Dunnett’s multiple comparisons test. ∗p≤0.05, ∗∗∗p≤0.001. (G) Boxplot showing frequencies of monocyte-derived macrophages out of all leukocytes in the non-ischemic left hemisphere of sham controls, and mice induced with MCAO who were either injected with PBS, 10M aMOG CD4/CD8 CAR T cells, 10M aMOG CD4 CAR T cells, or 10M BDNF-secreting aMOG CD4/CD8 CAR T cells immediately following MCAO. Statistical significance was determined using an ordinary one-way ANOVA with Dunnett’s multiple comparisons test. ∗∗∗∗p≤0.0001. (H) Boxplot showing frequencies of cDC2 cells out of all leukocytes in the non-ischemic left hemisphere of sham controls, and mice induced with MCAO who were either injected with PBS, 10M aMOG CD4/CD8 CAR T cells, 10M aMOG CD4 CAR T cells, or 10M BDNF-secreting aMOG CD4/CD8 CAR T cells immediately following MCAO. Statistical significance was determined using an ordinary one-way ANOVA with Dunnett’s multiple comparisons test. ∗∗∗∗p≤0.0001. (I) Boxplot showing frequencies of Xcl1+ CD8 T cells out of all leukocytes in the brain of sham controls, and mice induced with MCAO who were either injected with PBS, 10M aMOG CD4/CD8 CAR T cells, 10M aMOG CD4 CAR T cells, or 10M BDNF-secreting aMOG CD4/CD8 CAR T cells immediately following MCAO. Statistical significance was determined using an ordinary one-way ANOVA with Dunnett’s multiple comparisons test. ∗p≤0.05. (J) Boxplot showing frequencies of Xcl1+ CD4 T cells out of all leukocytes in the brain of sham controls, and mice induced with MCAO who were either injected with PBS, 10M aMOG CD4/CD8 CAR T cells, 10M aMOG CD4 CAR T cells, or 10M BDNF-secreting aMOG CD4/CD8 CAR T cells immediately following MCAO. Statistical significance was determined using an ordinary one-way ANOVA with Dunnett’s multiple comparisons test. ∗∗p≤0.01. (K) Boxplot showing frequencies of cDC1 cells out of all leukocytes in the brain of sham controls, and mice induced with MCAO who were either injected with PBS, 10M aMOG CD4/CD8 CAR T cells, 10M aMOG CD4 CAR T cells, or 10M BDNF-secreting aMOG CD4/CD8 CAR T cells immediately following MCAO. Statistical significance was determined using an ordinary one-way ANOVA with Dunnett’s multiple comparisons test. ∗p≤0.05. (L) H&E staining of a representative mouse brain 5 days post 60 minute occlusion of the MCA and adoptive transfer of 10 million BDNF-secreting CAR T cells. Large infarcts and ischemic foci seen in all sections.

**Figure S4.**
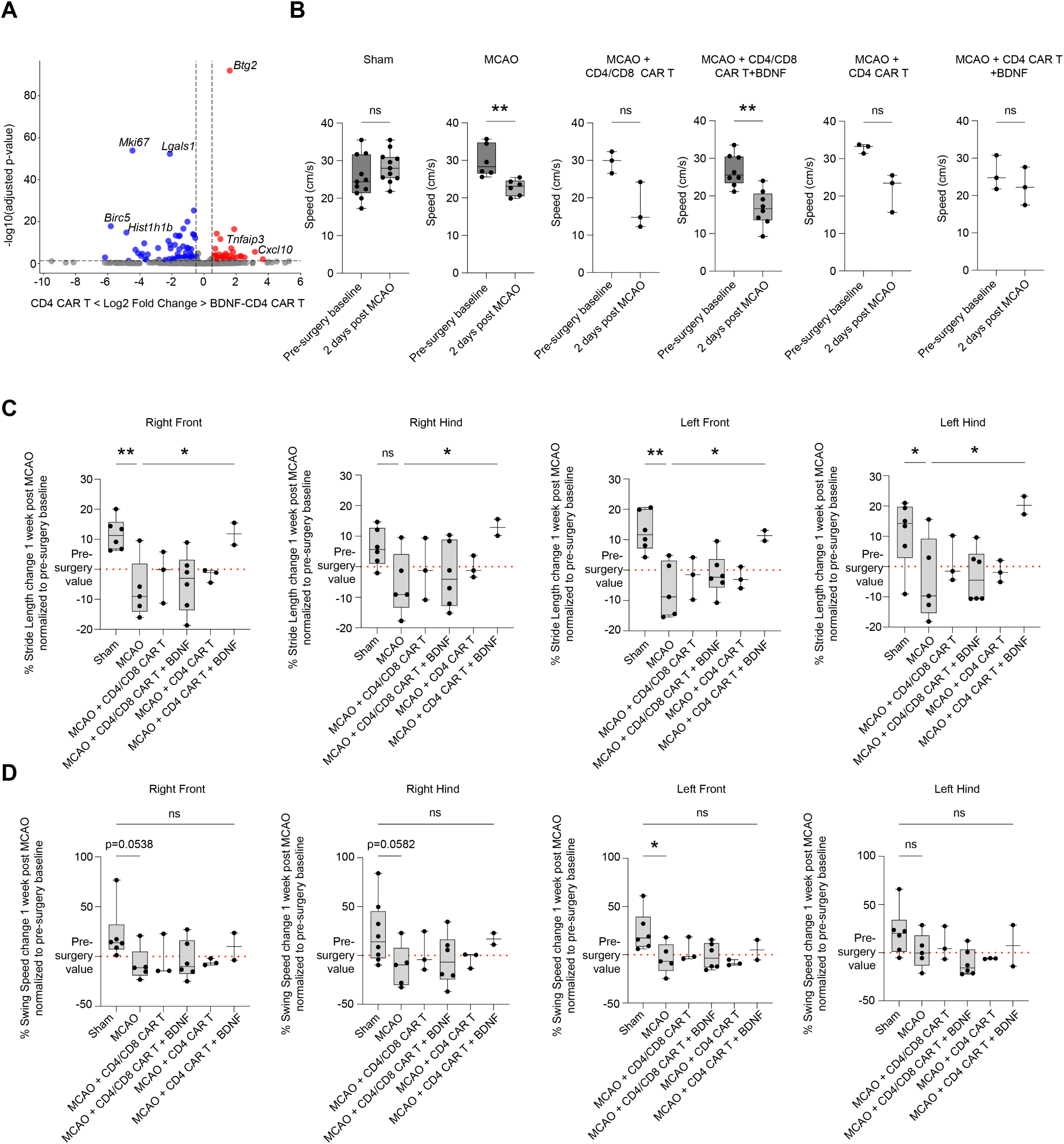
BDNF-secreting CD4 CAR T cells improve functional recovery after stroke, related to Figure 3. (A) Volcano plot of pseudo-bulk differential surface gene expression analysis between BDNF-CD4 CAR T cells and CD4 CAR T cells in brain of mice induced with MCAO. Top surface genes are highlighted and labeled as indicated. Differential gene expression has been tested using DESeq2. Dashed lines depict threshold for genes within p(adj) < 0.05. (B) Boxplots comparing speed (cm/s) pre-surgery and 2 days post MCAO surgery, and depicting sham controls and mice that underwent MCAO with gait deficits, who were either injected with PBS, 10M aMOG CD4/CD8 CAR T cells, 10M aMOG CD4 CAR T cells, 10M BDNF-secreting aMOG CD4/CD8 CAR T cells or 10M BDNF-secreting aMOG CD4 CAR T cells, immediately following MCAO. Statistical significance was determined using a paired t test. ∗∗p≤0.01. (C) Boxplot showing percent stride length change 1 week post MCAO surgery, across all paws, normalized to pre-surgery baselines, and depicting sham controls and mice that underwent MCAO with gait deficits, who were either injected with PBS, 10M aMOG CD4/CD8 CAR T cells, 10M aMOG CD4 CAR T cells, 10M BDNF-secreting aMOG CD4/CD8 CAR T cells or 10M BDNF-secreting aMOG CD4 CAR T cells, immediately following MCAO. Statistical significance was determined using an ordinary one-way ANOVA with Sidak’s multiple comparisons test. ∗p≤0.05, ∗∗p≤0.01. (D) Boxplot showing percent swing speed change 1 week post MCAO surgery, across all paws, normalized to pre-surgery baselines, and depicting sham controls and mice that underwent MCAO with gait deficits, who were either injected with PBS, 10M aMOG CD4/CD8 CAR T cells, 10M aMOG CD4 CAR T cells, 10M BDNF-secreting aMOG CD4/CD8 CAR T cells or 10M BDNF-secreting aMOG CD4 CAR T cells, immediately following MCAO. Statistical significance was determined using an ordinary one-way ANOVA with Sidak’s multiple comparisons test. ∗p≤0.05.

